# Comparative analysis of transposable elements in jellyfish and hydroid species (Cnidaria: Medusozoa)

**DOI:** 10.64898/2026.04.17.719288

**Authors:** Ayanna Mays, Feresa Cabrera, Aide Macias-Muñoz

**Affiliations:** Department of Ecology and Evolutionary Biology, University of California, Santa Cruz, CA 95064 U.S.A.

**Author notes:** **Corresponding authors:** Ayanna Mays,; Aide Macias-Muñoz.

**Keywords:** transposable elements, Cnidaria, Medusozoa, evolution, comparative genomics

## Abstract

**Background:** Transposable elements (TEs) are repetitive genetic elements that can jump to new loci causing genome expansions, structural rearrangements, and can, ultimately, propel the evolution of genomes. Despite their significance, the role of TEs in the evolution of genomes and phylogenetic groups remains largely understudied in early diverging lineages. Further, the extent to which TE content varies across species is still an open question. Medusozoa, a group within Cnidaria encompassing jellyfish and hydroids, exhibits an exceptional diversity of life history strategies, body plans, and physiological capabilities. These characteristics, along with its early-diverging phylogenetic position, establish Medusozoa as an ideal system for investigating the composition and evolutionary history of TEs within the group.

**Results:** We generated a custom repeat library built from annotations of 25 Medusozoan genomes and used it to characterize TEs, aiming to identify lineage-specific TE content and activity that may correlate with the diversity observed within the group. We found that repetitive element percentage and genome size varied considerably, with Hydrozoa exhibiting the most variation among classes in both respects. DNA transposons were the most prevalent TE classification in all but two genomes, averaging 28% of all genomes. Intra-genus comparisons revealed a surprising degree of differences in TE content. In the genus *Aurelia*, the expansion of a single DNA transposon superfamily accounted for much of the difference in repetitive element percentage between two species, whereas in the genus *Turritopsis*, a similar divergence resulted from the proliferation of multiple superfamilies. Interestingly, most genomes showed evidence of recent TE expansions, suggesting ongoing activity in many medusozoan species.

**Conclusion:** We present the first comparative analysis of TEs across all medusozoan classes. Our results reveal class-specific TE dynamics and highlight cases of TE proliferations as lineages diverge. This research provides data on TE activity and diversity that can be used as a resource for future study and fills important gaps in our understanding of TEs in early diverging animal lineages.

## Background

Transposable elements (TEs) are repetitive sequences of DNA that can insert themselves into new genomic loci, rapidly generating genetic variation. TEs are grouped into two classes based on their transposition mechanism (1–3). Class I Retrotransposons mobilize via a “copy and paste” mechanism involving an RNA intermediary. Class I elements are further divided into LTR (long terminal repeat) and non-LTR (non-long terminal repeat) elements, which include LINEs (long interspersed nuclear elements) and SINEs (short interspersed nuclear elements). Class II DNA Transposons move via a “cut and paste” mechanism and consist of TIR (terminally inverted repeat), Helitron (rolling circle), and Maverick elements that are further organized into superfamilies including hAT and Tc1/mariner. Together with other repetitive elements like simple, satellite, and interspersed repeats, TEs can expand genome size, reorganize genome structure, and importantly, lay the groundwork for genomic innovation by introducing novel genotypes to populations. These changes provide natural selection an opportunity to act, propelling evolution within genomes and, ultimately, driving diversification across lineages (4,5)

The discovery of TE-driven genomic changes has shifted our understanding of genomes from that of static units that evolve at the rate of point mutations and recombination alone. Rather, they are dynamic entities that can undergo rapid changes that express phenotypically to shape organismal evolution, morphology, and adaptive traits. Factors that shift a genome away from its ancestral state range from smaller scale processes like gene gains and losses to larger scale events including, chromosomal rearrangements and changes in genome size. TEs often facilitate gene gain through gene duplications driven by mechanisms like gene retrotransposition and non-allelic homologous recombination (NAHR) (6–9). The result is two copies of a gene that follow different evolutionary trajectories where one copy may acquire an entirely new function (10,11). In aphids, for instance, TEs have contributed to the evolution of expanded gene families related to sensory functions. Over time, the compounding effects of TEs continually pushes a genome further from its ancestral state. Organisms have, however, evolved defense mechanisms against TE activity as their movement can result in genomic instability. The PIWI-piRNA pathway, conserved among metazoans, functions to repress TE movement (12). DNA methylation, a mechanism of gene regulation, also preferentially silences TEs (13,14). This suppression can eventually render TEs inactive or “domesticated” and can then be co-opted for novel functions (15,16). For instance, many transcription factors across Metazoa are derived from domesticated TEs (17). The relationship between TEs and their genomic habitats is inextricably linked with both parties evolving mechanisms and functions in response to one another. As such, integrating TEs into comparative analyses is essential for understanding not only genome evolution, but also broader patterns of organismal evolution across taxa.

Cnidaria, the sister group to Bilateria, arose approximately 741 million years ago and serves as a valuable reference point for early animal evolution (18). The phylum encompasses two major clades: Anthozoa (corals and sea anemones) and Medusozoa (jellyfish and hydroids). Within Medusozoa, four classes are recognized: Cubozoa, Staurozoa, Hydrozoa, and Scyphozoa (19,20). Medusozoa is a particularly useful system of genome evolution because its species display a wide range of body plans, reproductive strategies, and life history traits. The medusa stage, for instance, is a life cycle stage that has evolved independently and been lost multiple times (21–23). Additionally, Medusozoa includes species with unique biological capabilities. *Turritopsis dohrnii*, for example, is renowned for its ability to undergo life stage reversal under stress, a phenomenon often described as biological immortality (24). The phylogenetic position and extensive plasticity of body plans and life history strategies in Medusozoa make it an ideal group to study the evolutionary forces that have shaped its diversity. Despite this, TE activity is underexplored here as well as in other early diverging animal lineages. Previous work has begun to identify and characterize TEs in some Medusozoan species at the individual level (25–28). In *Hydra vulgaris*, a lineage specific expansion of the LINE/CR1 superfamily occurring after the separation of brown and green hydra was discovered (29). The expansion of the DNA transposon, Helitron, in *Hydractinia symbiolongicarpus* has also been identified (30). While these studies begin to fill in gaps about the evolution of TEs in select Medusozoan species, the role that they have played in broadly shaping the group’s evolution remains elusive.

Here, we characterized TEs in 25 Medusozoan species and reconstructed their activity over evolutionary time using Kimura divergence analysis with the goal of finding patterns in TE content and activity that may explain the wide phenotypic variation observed in the group. Results from our phylogenetic reconstruction resolved inter-class relationships within Medusozoa. We examined the relationship between genome size and repetitive element content and quantified the relative abundance of TE families across genomes. Using repeat landscape graphs, we visualized periods of repetitive element proliferation and compared these patterns across both closely and distantly related species. We found remarkable diversity in TE content and activity across medusozoan genomes. In addition, we found that the strength of the relationship between genome size and repetitive element percentage differed across medusozoan classes. In some instances, we were able to pinpoint specific TE families that proliferated after the divergence of closely related species. Altogether, these results offer new insights into medusozoan genome evolution and establish a framework for future comparative studies of TEs in early diverging animal lineages.

## Methods

### Genome Acquisition

We downloaded publicly available genomes for 25 Medusozoan and one Anthozoan species from NCBI GenBank to UCSC’s high-performance computing cluster using the wget command. The genomes used in this study along with relevant genome quality metrics can be found in Supplemental Table S1 (Table S1).

### Phylogenetic Analysis of Cnidarian Mitogenomes

#### Mitochondrial Data Acquisition and Recovery

Mitogenomic data were obtained from a combination of public repositories and *de novo* assemblies. Specifically, nucleotide sequences of 21 previously published cnidarian mitochondrial genomes were retrieved from GenBank (Table S1). For additional taxa where mitogenome data were unavailable, we downloaded corresponding genomic Sequencing Read Archive (SRA) datasets and performed *de novo* assembly using GetOrganelle 1 (v1.7.7.1). To ensure consistent gene boundary prediction across all samples, every mitochondrial genome, both published and newly assembled, was uniformly re-annotated using MitoFinder 2 (v1.4.2). From the standardized annotations, we extracted the nucleotide sequences of 13 protein-coding genes and two ribosomal RNA (rRNA) genes for phylogenetic analysis. The protein-coding genes included ATP synthase F0 subunit 6 and 8 (*atp6* and *atp8*), cytochrome b (*cob*), cytochrome c oxidase subunits I–III (*cox1– 3*), NADH dehydrogenase subunits 1–6 and 4L (*nad1–6* and *nad4L*), and the large and small ribosomal RNA subunits (16S *rrnL* and 12S rrnS).

#### Sequence Alignment and Dataset Construction

The 13 protein-coding genes (*atp8, atp6, cox1, cox2, cox3, cob, nad1, nad2, nad3, nad4, nad4L, nad5, nad6*) and 2 ribosomal RNA genes (*rrnL* and *rrnS*) were aligned separately using MAFFT 7.471 with the L-INS-i algorithm. Alignment trimming was performed using Gblocks (31), with gaps allowed for all positions and a flanking position conservation threshold of 85% of the total number of sequences. To determine the best-fitting evolutionary models for each gene partition, we used ModelTest-NG 3 (v0.1.7). The resulting individual gene alignments were then concatenated into a single partitioned matrix for phylogeny reconstruction. The final concatenated dataset consisted of 374 nucleotide sequences spanning 24 distinct taxa. Phylogenetic reconstruction was performed using Anthozoan (*Nematostella vectensis and Haliclona simulans*) as the outgroup.

#### Phylogenetic Inference

Maximum likelihood (ML) analysis was conducted on the concatenated amino acid dataset using RAxML-NG (v0.8.1). The analysis was partitioned by gene, with the evolutionary models applied as determined by ModelTest-NG. The tree search employed 50 random and 50 maximum parsimony starting trees. Nodal support was assessed with 1000 bootstrap pseudoreplicates. Bayesian inference (BI) was performed using MrBayes (v3.2.7a) under the default settings.

### TE characterization

We generated a custom Medusozoa repeat library based on previously described methods with some modifications (32). Briefly, we used RepeatModeler v2.0.5 (33) to create custom repetitive element libraries for each genome with default settings (BuildDatabase -name species_name_db species_name.fna). We merged the resulting libraries using the cat function, removed redundancies using CD-hit (80-80-80 rule), and re-annotated sequences identified as “Unknown” with DeepTE (34). The resulting Medusozoa repeat library was used as input in the RepeatMasker function to identify and quantify the percentage of repetitive elements in each genome (RepeatMasker -lib medusozoa_rep_lib.fa species_name.fna -a).

### Kimura divergence and Repeat Landscape analysis

The Kimura substitution analysis calculates a sequence’s divergence from a determined consensus sequence. With the understanding that TEs accumulate mutations over time and the assumption that this accumulation is constant, how diverged a TE sequence is can be used as a proxy for age (35). We used the calcdiv function in RepeatModeler with the .align file generated from RepeatMasker to perform Kimura two-parameter analysis on each repetitive element identified (calcDivergenceFromAlign.pl -s species_name.fna.divsum species_name.fna.align). Then, following previously described analysis, we created RepeatLandscape graphs using resulting .divsum files in RStudio to visualize expansions and reductions of TE subfamilies over evolutionary time (36).

## Results

### Newly assembled mitogenomes and phylogenetic relationships in Medusozoa

To augment the available molecular data for Medusozoa, we assembled mitochondrial genomes from publicly available Sequence Read Archive (SRA) datasets for species where no complete mitogenome was previously published. This effort yielded two novel, linear mitochondrial genomes: one from the hydrozoan *Bougainvillea muscus* and another from the staurozoan *Calvadosia cruxmelitensis*. We additionally extracted additional gene markers from *Hydra vulgaris* (strain AEP) (ATP synthase subunits *atp6, atp8*; cytochrome c oxidase subunits *cox1, cox2, cox3*; cytochrome b *cytb*; NADH dehydrogenase subunits *nd1–nd6, nd4L*; and ribosomal RNA genes *rrnL, rrnS*) and *Chironex yamaguchii* (*atp6, atp8, cox2, cox3, nd1, nd3, nd4, nd4L*), expanding taxon sampling beyond currently published datasets. The assembly and structure of these genomes are detailed in Fig S1. These new data fill important taxonomic gaps, particularly within the understudied Staurozoa.

Using these newly assembled mitogenomes and some extracted from our species of interest, we reconstructed a phylogeny to resolve the inter-class relationships within Medusozoa. Our analysis included a dataset of 25 mitochondrial genomes, selected to represent all four medusozoan classes Hydrozoa, Scyphozoa, Cubozoa, and Staurozoa with an anthozoan used as the outgroup (see Methods, Table S2). Our phylogenetic inference (Fig S2) recovered a topology that supports the monophyly of each class, consistent with prior mitochondrial-based studies (20,37). However, it also reveals a distinct hypothesis for inter-class relationships. Contrary to the prevailing phylogenomic framework that groups Cubozoa and Scyphozoa as sister classes with Staurozoa as their sister group (37), or recover alternative topologies (38), our mitochondrial phylogeny recovered Staurozoa as the sister lineage to Cubozoa. This relationship was strongly supported by the placement of the newly assembled mitogenome of *Calvadosia cruxmelitensis*, which, together with the published mitogenome of *Haliclystus octoradiatus*, forms a staurozoan clade that is sister to Cubozoa (represented here by *Morbakka virulenta* and *Chironex yamaguchii*). This result suggests that mitochondrial genomic data may retain a divergent phylogenetic signal regarding the relationships among Scyphozoa, Cubozoa, and Staurozoa, underscoring the need for further investigation into the evolutionary history of these lineages.

### Relationship between genome size and repetitive elements

To investigate the relationship between genome size and repetitive elements, we compared genome size and repetitive element percentage across all genomes. Genome size ranged from 209 Mb to 3327 Mb, with repetitive elements comprising 24% to 78% of genomes (Fig 1B, Table S3). Hydrozoa exhibited dramatically more genome size variation than other classes. The range of genome sizes in Hydrozoa (3095 Mb) exceeded a combined range of Scyphozoa, Cubozoa, and Staurozoa by more than fourfold (742 Mb) and all genomes greater than 500 Mb were hydrozoan, except for two species: the scyphozoan *Aurelia coerulea* and the cubozoan *Morbakka virulenta*. Notably, two sizable genomes over 3 Gb were observed in Hydrozoa: *Physalia physalis* and *Millepora dichotoma*, representing some of the largest cnidarian genomes sequenced to date (Fig 1B, Table S2). Repetitive element percentages also varied substantially across classes. The lowest percentage observed was in the scyphozoan genome *Nemopilema nomurai* (24%), while the highest occurred in hydrozoans; two strains of *H. vulgaris* reached 77.11% in strain AEP and 78% in strain 105.

**Figure 1.**
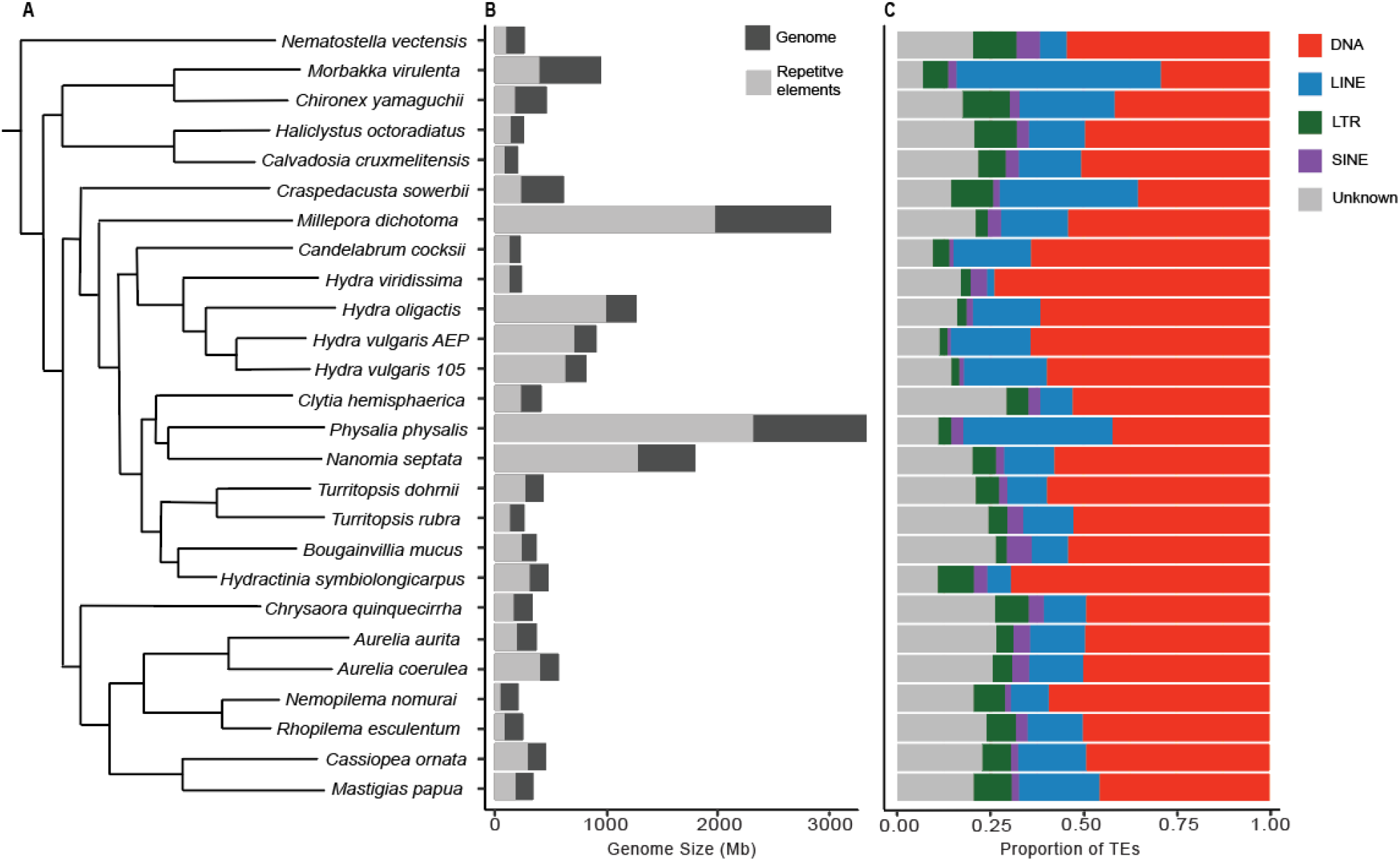
Repetitive element percentage and transposable element (TE) content in Medusozoan genomes. A. Cladogram based on phylogenetic reconstruction of 25 Medusozoan and 1 Anthozoan species (Fig S1). B. Amount of repetitive elements (light gray) in relation to genome size (dark gray) in Megabases (Mb). C. Proportion of TE classes in relation to total classified TEs.

Within genus comparisons revealed substantial differences in repetitive element percentage. For instance, in the genus *Aurelia*, repetitive elements made up 70% of the *A. coerulea* genome but only 52% of that of *A. aurita*, an 18% difference. In the genus *Turritopsis*, the difference was less extensive but still substantial by 11% between *T. dohrnii* (62%) and *T. rubra* (51%). These genus-level differences suggest that TE content can shift on relatively short evolutionary timescales. Intraspecific variation in repetitive element percentage was minimal: *Hydra vulgaris* strains AEP and 105 differed by less than 1% (Table S3). The drastic difference in TE percentage between congeners compared to nearly identical profiles of conspecific strains suggests that TE accumulation is not solely gradual in Medusozoa.

The relationship between genome size and repetitive element percentage is generally linear and positive, such that larger genomes typically contain more repetitive elements (39–41). We performed linear regression models to test whether this expectation holds in Medusozoa and found that many genomes deviate from this relationship (Fig 2). Additionally, the strength of this relationship differs among classes. Across all genomes, we obtained a Pearson correlation coefficient of r^2^=0.44, displaying a moderate, positive correlation (Figure 2, Table S3). When we performed this analysis on Scyphozoa alone, one of two classes with sufficient genome sampling, we obtained a strong, positive correlation (r^2^=0.95) (Table S3). In contrast, Hydrozoa alone showed a weak, positive correlation (r^2^=0.39), indicating that hydrozoan genomes follow this expectation more loosely, shifting the result of all medusozoan genomes from a strong correlation of genome size and repetitive element percentage to a moderate one. Indeed, when hydrozoan genomes were excluded from analysis, we obtained a stronger correlation (r^2^=0.72). These results indicate that the relationship between genome size and percent of repetitive elements vary among classes, possibly reflecting class-specific evolutionary pressures on TE accumulation.

**Figure 2.**
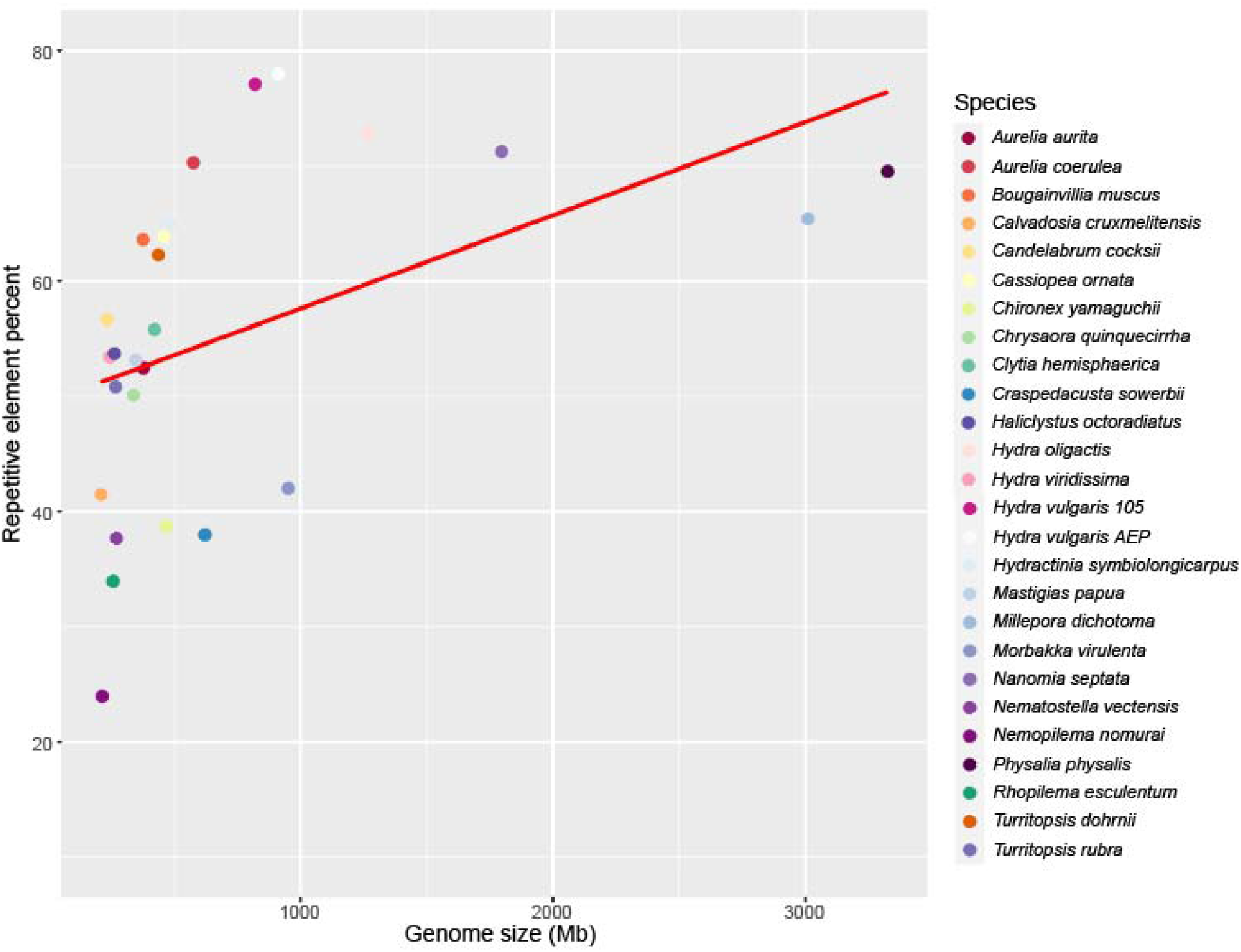
Genome size by repetitive element percentage shows a statistically significant positive correlation (r^2^=0.44). Each circle is color coded by species and the red line represents a fitted linear regression.

### Transposable element content and diversity

To understand differences in repetitive element content among genomes, we quantified the percentage of major TE classifications present including DNA transposons, LINE, SINE and LTR elements. DNA transposons made up an average ∼28% of repetitive elements classified across genomes and were the most dominant TE classification in all but two species, *Morbakka virulenta* and *Craspedacusta sowerbii*, where LINEs predominated instead (Fig 1C). LINEs were the second most dominant, excluding two species, averaging ∼9%, followed by LTR elements averaging ∼3% and SINEs averaging ∼2% (Table S4). Broadly, while genome size and TE percentage greatly differed, the relative proportions of TE classes differed to a lesser degree. Even with this, all major TE classifications were present in Medusozoa.

The 9 representative scyphozoan genomes exhibited generally consistent proportions of TE classifications. DNA transposons were the most prevalent in all scyphozoan genomes, averaging ∼24%. In *Chrysaora quinquecirrha*, a species recovered as a sister to all other scyphozoans, DNA transposons were the most dominant (23%) followed by LINEs (5.33%), LTR (4.26%), and SINEs (1.99%). *Nemopilema nomurai* was composed of 13.35% DNA transposons, 2.3% LINEs, 1.9% LTR, and 0.39% SINEs. Its sister species, *Rhopilema esculentum*, shows slightly higher proportions: 16.45% DNA transposons, 4.89% LINEs, 2.60% LTR, and 1.08% SINEs. As mentioned before, we observed a difference in repetitive element percentage in the genus *Aurelia*. We found that DNA transposons were largely responsible for this difference evidenced by a 9% increase in *A. coerulea* compared to *A. aurita*. The genome of *A. coerulea* contains 34.48% DNA transposons, 10.08% LINEs, 3.31% SINEs, and 3.58% LTR elements. In contrast, *A. aurita* has 25.46% DNA transposons, 7.63% LINEs, 2.36% LTR, and 2.39% SINEs (Table S4).

Comparing the TE composition across hydrozoan genomes revealed class-specific trends, with greater variation in TE classification proportions than other classes. Consistent with Scyphozoa, DNA transposons were the most prevalent except in *Craspedacusta sowerbii*, averaging 36.59% of genomes, with *Hydra viridissima* and *Hydractinia symbiolongicarpus* showing the highest proportions (Fig 1C). *Craspedacusta sowerbii* shows a proportionally large presence of LINEs and the smallest percentage of DNA transposons (12.13%) compared to other hydrozoans (28.22%-46.46%) (Fig 1C, Table S4). In the *Hydra* genus, LINEs are nearly absent in *H. viridissima* (2%) but are expanded in other *Hydra* species (13-16%), corroborating previous reports of an independent LINE expansion in brown hydra (29) (Fig 1C). In *Turritopsis*, DNA transposons account for most of the difference in repetitive element content between *T. dohrnii* and *T. rubra. T. dohrnii* is composed of 35.35% DNA compared to 24.82% in *T. rubra*, an 11% difference (Table S4). The differential expansions of DNA transposons in *Hydra, Turritopsis*, and *Aurelia* indicate multiple independent expansions across Medusozoa. The expansion of LINE in *Hydra* is not recapitulated elsewhere, suggesting this is a unique event restricted to brown hydra.

Differences in TE content between the two representative staurozoan species, *Calvadosia cruxmelitensis* and *Haliclystus octoradiatus*, were minimal. DNA transposons differed by ∼5% between the two species, while LINEs, SINEs, and LTR elements differed by 0.94%, 0.27%, and 2.86% respectively. On the contrary, there were substantial differences in TE content between the cubozoan species *Morbakka virulenta* and *Chironex yamaguchii*. Most substantially, LINE percentage in the respective genomes differed by 12%. DNA transposons differed by 3.95%, while other TE classifications were more consistent between the two. Repetitive element data for all genomes can be found in Table S4.

### Evolutionary trajectories of TEs in Medusozoa

The natural mutation rate of TEs after insertion allows us to estimate their relative age based on sequence divergence. More recently inserted TEs cluster to the left of the x-axis while older insertions cluster towards the right. Overall, we observed substantial variation in TE activity at both the class and genus level. In some cases, single superfamily expansions contributed to TE peaks, while other peaks were composed of multiple different types.

Cubozoan genomes, *Morbakka virulenta* and *Chironex yamaguchii*, showed strikingly different profiles of TE activity over time. *Morbakka virulenta* had at least 3 distinct expansion periods, all of which were dominated by several LINE superfamilies, the most abundant of which were LINE/RTE and LINE/Dong elements. In contrast, its sister species, *Chironex yamaguchii* showed a single ancient expansion peak, composed primarily of DNA transposons, followed by a recent contraction across all superfamilies. As such, it is unlikely that *Chironex yamaguchii* contains currently active TEs.

We included two staurozoan genomes, but a repeat landscape for one, *Haliclystus octoradiatus*, is not reported as results did not match expectations and we consider them unreliable for interpretation (Figure S3). The other, *Calvadosia cruxmelitensis*, exhibits a continuous expansion of TEs with a recent peak (substitution levels 0-5). Here, DNA transposon superfamilies DNA/TcMar and DNA/hAT primarily drive the steady increase in DNA transposons, which are only recently contracted. MITE elements and LINE superfamilies like CR1 and RTE have also contributed to TE accumulation in this genome. See data availability for access to interactive versions of Kimura substitution graphs.

TE activity in Hydrozoan genomes also varied widely. *Craspedacusta sowerbii* displays two distinct expansion periods, each driven by different TE superfamilies. LINE/L2 were primarily responsible for the more ancient peak, while DNA transposons DNA/hAT and DNA/TcMar expanded more recently. Similarly, *Millepora dichotoma* also shows an ancient expansion of LINE/L2 elements and a more recent expansion of DNA/hAT and Tc/Mar, though to a greater overall extent. Though these expansion patterns are unique to these two species, their divergence time suggests they arose independently. TE activity in *Hydra* reflects lineage-specific patterns. As previously noted, LINE/CR1 and LINE/L2 elements were expanded in brown hydra (*H. vulgaris* and *H. oligactis*) species but absent in green hydra (*H. viridissima*) (Fig 1C, Fig 3). What could not be discerned from bulk TE percentages, however, is the recent expansion of SINE in *H. viridissima*. Though there is just over a 1% difference in overall SINE elements between brown (0.74-0.97%) and green (2.13%) hydras, much of this difference occurred in a very recent expansion, suggesting SINEs may be currently active in green hydra alone. The evolutionary history of TEs in *Turritopsis* highlights species-specific characteristics. The difference in DNA transposon percentage between *T. dohrnii* and *T. rubra* is notable given their close relatedness. Unlike other cases where a single superfamily is responsible for a TE class expansion, several DNA superfamilies contributed to the difference seen between these two species, including DNA/TcMar, DNA/hAT, and DNA/Maverick with the former being the most differentially expanded. Expansion peaks in other genomes, like *Bougainvillia muscus* and *Hydractinia symbiolongicarpus*, were also dominated by DNA/hAT, DNA/Mutator, and DNA/TcMar.

**Figure 3.**
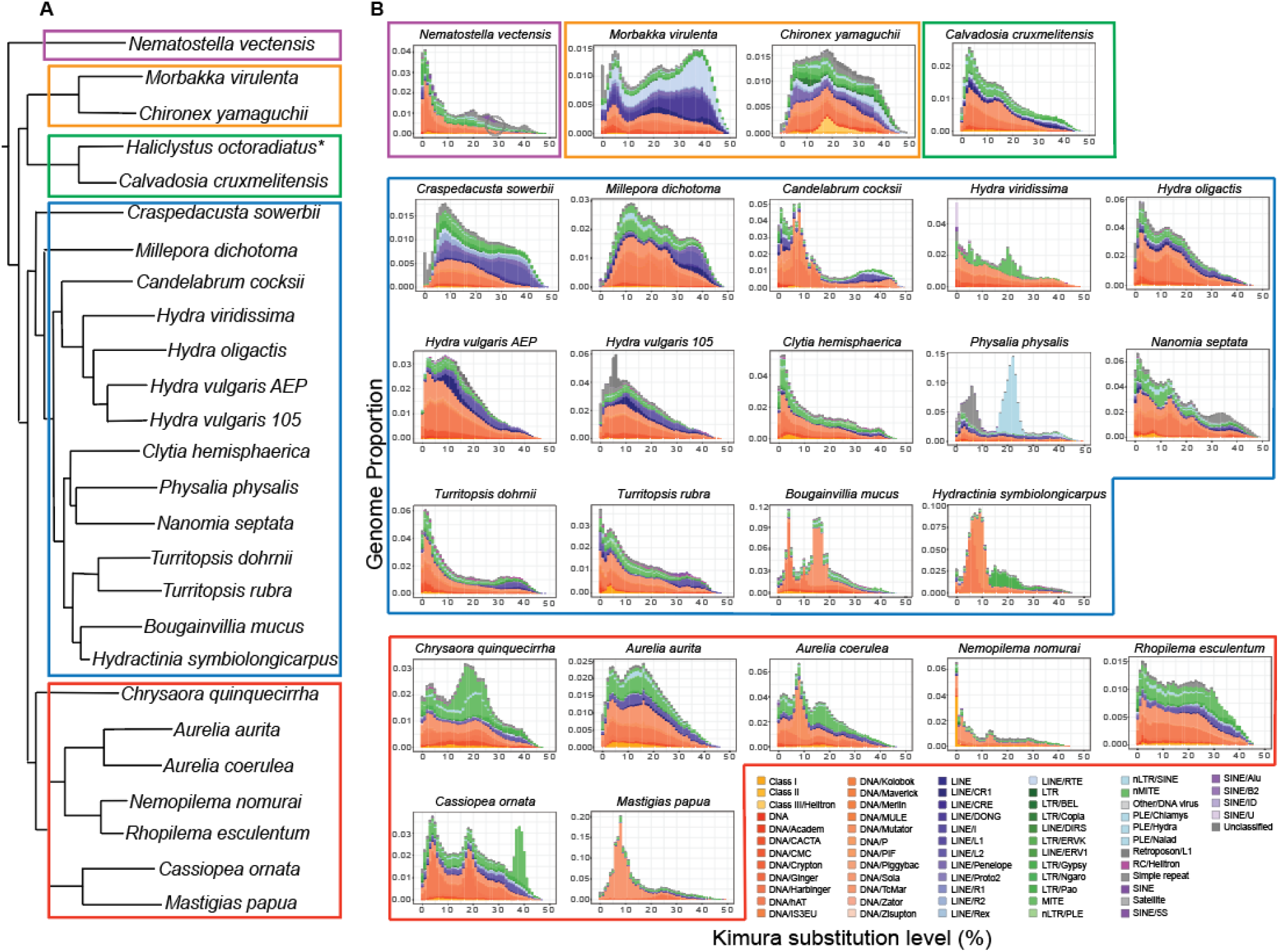
Transposable element expansions over relative evolutionary time. A. Cladogram of phylogenetic relationships between species. Boxes denote cnidarian classes: purple - Anthozoa, orange - Cubozoa, green - Staurozoa, blue - Hydrozoa, red - Scyphozoa. B. Repeat landscape graphs of focal species with boxes mirroring the cladogram. Peaks convey bursts of TE activity over relative time. The legend (lower right) labels TE superfamilies identified by RepeatMasker. Note that the scale of the y-axis varies. \**Haliclystus octoradiatus* included in the supplement (Fig S3).

Scyphozoan genomes exhibit striking variation in TE activity. Some species show multiple expansion peaks, some are dominated by only a few superfamilies, and others display greater diversity (Fig 3). *Chrysaora quinquecirrha* has 2 expansion peaks and shows a notable expansion of MITE elements not seen in other scyphozoan genomes. *Cassiopea ornata* exhibits three distinct peaks, with the most ancient being primarily MITE elements and the two more recent ones composed of DNA/Harbinger and DNA/hAT. The genomes of *Mastigias papua* and *Nemopilema nomurai* show considerably less diversity than other scyphozoans, with a single DNA transposon, DNA/TcMar, contributing 89% to a peak expansion. In *Aurelia, A. coerulea* displays an expansion of DNA/mutator that is nearly absent in *A. aurita*. This expansion likely accounts for the previously noted difference in repetitive element content between the two species.

## Discussion

Transposable elements are prevalent components of eukaryotic genomes. Their movement can generate genetic mutations, facilitate gene duplications, and introduce novel sequences that can be co-opted to form new genes or regulatory elements. As such, they contribute substrate for genomic innovation and are associated with speciation events, altogether affecting the evolutionary trajectory of both individual genomes and entire lineages (4,5). A comparative understanding of TE content and activity in the early-diverging Medusozoa offers a critical window into the role that TEs have played in shaping animal genome evolution. Previous genome assemblies and small-scale comparative studies have begun to reveal the genomic changes accompanying species divergence in this group. For instance, one study identified gene family expansions and contractions in the giant Nomura’s jellyfish (*N. nomurai*) relative to the common ancestor of genera *Nemopilema* and *Aurelia* (42). Another case finds the rate of evolution of a scyphozoan genome to be slower than that of a hydrozoan (43). While these and other efforts (44–46) have advanced our understanding medusozoan genome evolution, few (47) comparatively analyze repetitive elements and explore their relevance in genome evolution. The findings of this study fill this gap and reveal the unique characteristics of medusozoan classes, genomes of notable size and TE content, and TE expansions that may be of interest for future studies.

### Phylogenetic groupings in Medusozoa

Our mitogenome phylogenetic analysis served to clarify relationships and contribute newly constructed mitogenomes in underrepresented classes. Our analysis robustly recovers Staurozoa as sister to Cubozoa, revealing a strong signal for a close evolutionary relationship between these two medusozoan lineages. This result is particularly notable given that Staurozoa has historically been underrepresented in genomic datasets, limiting resolution of its evolutionary placement. By incorporating mitogenomic data that improve representation of this lineage, our analysis provides clearer insight into its position within Medusozoa and helps resolve a long-standing gap in cnidarian phylogeny.

The Staurozoa-Cubozoa sister relationship recovered here finds a notable parallel in recent morphological and taxonomic revisions. Work reorganizing Rhopaliophora (Scyphozoa, Cubozoa and Staurozoa) based on early life cycle characters (48,49) has proposed the inclusion of both Cubozoa and Staurozoa within an expanded Scyphozoa. While our data do not support a single, expanded Scyphozoa clade, they echo the proposed close evolutionary link between Staurozoa and Cubozoa suggested by shared developmental traits. The congruence between our mitochondrial signal and this independent phenotypic hypothesis underscores the need to integrate diverse data types to resolve deep medusozoan relationships. However, the phylogenetic position of Staurozoa remains contentious. A large-scale mitochondrial study of 266 cnidarian genomes recovered a contrasting topology, placing Staurozoa closer to Anthozoa (38). Importantly, their sampling, like earlier mitochondrial-based studies (37) included only a single staurozoan genus (*Haliclystus*), whereas our dataset incorporates two staurozoan genera (*Haliclystus* and *Calvadosia*), improving lineage representation within the group. This difference in sampling breadth likely contributes to the conflicting topologies: our Medusozoan-focused phylogeny captures finer-scale relationships that may be diluted in broader, phylum-level analysis. Taken together, these results demonstrate that increasing genomic representation in historically understudied lineages like Staurozoa substantially improves phylogenetic resolution.

### Increased TE diversity compared to other cnidarian groups

TE diversity has been well studied in other animal lineages, particularly vertebrates (39,50–53). In early diverging groups like Cnidaria, however, the extent of TE diversity is poorly understood. Recent work in Anthozoa, the sister group to Medusozoa, has begun to address this gap, and comparisons with these studies reveal striking differences between the two cnidarian clades. An analysis in the order Actiniaria, for instance, revealed considerably less TE landscape diversity with all species analyzed having one recent TE expansion (54). Similarly, in the repeat landscapes of Zoantharia, another Anthozoan order, landscapes are visually congruent with little variation in repetitive elements and expansion peaks (55). Conversely, here we identified a large degree of landscape variation within orders. For instance, in Anthoathecata (the genus *Hydra, M. dichotoma, P. physalis, C. cocksii, H. symbiolongicarpus*, and *B. mucus*), species display one, two, or three expansion peaks comprised of varying repetitive elements (Figure 3). The same is seen in another medusozoan order represented here, Rhizostomae (*N. nomurai, R. esculentum, C. ornata*, and *M. papua*), where species display two, two, three, and one expansion peaks respectively and are considerably different in repetitive element makeup (Figure 3). The seemingly increased level of TE diversity in medusozoan orders indicates that Medusozoa has experienced particular evolutionary pressures that drive TE activity to change rapidly and dramatically. In particular, the complexity of body plans compared to other cnidarian groups, the evolution of complex traits, such as the independent evolution of eyes, and the diverse habitats that they inhabit may be linked to this genomic observation.

### Repetitive elements and genome size relationship differs between classes

Repetitive elements are major drivers of genome variation and a positive correlation between TE content and genome size is often assumed (39–41). Our results broadly support this relationship across Medusozoa, but the strength of this correlation is not uniform and varies across classes. Though Medusozoa displays a high degree of TE diversity overall, Hydrozoa stands apart from other classes as having the most variation in genome sizes and the weakest correlation between genome size and repetitive element percentage (Supplementary table S3). Distinct from other classes, hydrozoan species have secondarily lost the medusa stage repeatedly (21–23), have invaded freshwater habitats (*Hydra, C. sowerbi*i) (56–59), and display a wide range of reproductive modes. These unique features of Hydrozoa and the variation of genome size and repetitive element percentage may be causally linked. Studies into the evolutionary history of genomes that fall far outside the predicted linear model will elucidate why some species have more or less repetitive elements than expected (Figure 2).

In contrast to Hydrozoa, Scyphozoa displays a strong and predictable TE-genome size relationship that aligns with canonical expectations and has been found in similar studies (47). Between these poles, Cubozoa and Staurozoa suggest additional modes of evolution, though both remain sparsely sampled. Cubozoa shows evidence of lineage-specific trajectories, where TE dynamics diverge even among close relatives, emphasizing that genome evolution can operate at fine phylogenetic scales. Staurozoa, in contrast, exhibits genomic constraint, with minimal variation in TE content and composition. Both patterns must be interpreted cautiously given limited sampling (60,61). This shift toward a lineage-dependent view of TE dynamics highlights a critical need for denser taxonomic sampling, especially in Cubozoa and Staurozoa, and for comparative approaches that move beyond one-size-fits-all models.

### TE are dynamic in Medusozoa

We compared TE evolution in closely related species to shed light on the degree and speed to which TEs evolve over relatively short timescales. Species in *Aurelia* have highly divergent protein sequences with their degree of difference being greater than that between mice and rats (44). The present study found that repetitive elements in *A. aurita* and *A. coerulea* also differed substantially and the DNA transposon superfamily, DNA/Mutator, is expanded in *A. coerulea* alone, indicating that this expansion occurred after their divergence approximately 90 mya (Figure 3, (62). In another instance, we compared the TE profiles of species in *Turritopsis* and found that, unlike Aurelia, multiple DNA families including DNA/CACTA, DNA/TcMar, and MITE elements drove differences between *T. dohrnii* and *T. rubra* genomes. *T. dohrnii* has the unique ability to undergo life cycle reversal in response to damage or stress, making it a functionally immortal organism. Whether and to what degree other species in the genus, like *T. rubra*, have this ability is not fully resolved (46,63–65). Studies have, however, delved into differences in genomic architecture between the two species (46). The proliferation of TEs between these two species may shed light on the underlying genomic features and evolutionary history that have given rise to the unique abilities seen in this genus. Broadly, these results present a basis for the role that TEs may have played in the diversification of lineages.

While we have demonstrated that TEs can proliferate rapidly in the genomes of closely related species, whether they stabilize at the species level remains an open question. Two strains of *H. vulgaris* had nearly identical TE profiles, with total content differing by just less than a percent. There were, however, slight differences best exemplified in the repeat landscape of *H. vulgaris* 105 which has an independent expansion of simple repeat elements not seen in *H. vulgaris* AEP. The very recent divergence time of these strains (calculated to be ∼16 mya) begs questions about drivers of TE turnover and proliferation aside from evolutionary time (29). Within species TE variation has been recorded in the green hydra, *H. viridissima*, where one individual contained functional copies of a DNA element and the other did not (26) showing that TE content can differ from individual to individual. Taken together, we show evidence of TE dynamism at the genus and species level and suggest that TE evolution is complex and cannot be attributed to divergence time alone.

As genomes from diverse taxa become increasingly available, we are presented with the opportunity to further our understanding of underexplored features of evolution like transposable elements. Comprehensive understanding of TEs has the potential to shed light on diversification events, their contribution to the development of novel traits, and the roles that they may now play in organismal function. Utilizing early diverging systems presents even greater potential to elucidate evolutionary processes relevant to organisms across the Tree of Life.

## Conclusion

This study provides the first broad-scale comparative analysis of transposable elements across Medusozoa offering new insight into how these genomic elements have shaped the evolutionary history of this early-diverging lineage. TE dynamics in Medusozoa are lineage-specific, varying in scale and timing across conspecific strains, congeneric species, and whole classes to produce distinct genomic strategies rather than a single evolutionary pattern. The association between TE diversity and phenotypic innovation, particularly in Hydrozoa, points to TEs as potential engines of evolutionary novelty. Unlocking this potential will require denser sampling, functional studies, and comparative frameworks that move beyond one-size-fits-all approaches. Altogether, this study reveals rapid turnover of TEs, highlights cases of TE proliferations as lineages diverge, and establishes Medusozoa as a powerful system for understanding how transposable elements shape genome evolution.

## Supplemental files

Figure S1. Mitochondrial genome data

Figure S2. Phylogenetic reconstruction

Figure S3. *Haliclystus octoradiatus* repeat landscape

Table S1. Genome accession and quality metrics

Table S2. Mitochondrial genome accession

Table S3. Genome size and repetitive element percentage

Table S4. Pearson correlation statistics

Table S5-S31. RepeatMasker tables

## Declarations

### Ethics approval and consent to participate

Not applicable

### Consent for publication

Not applicable

### Availability of data and materials

All sequencing data used for this manuscript was obtained from publicly available genomes detailed in Supplementary Table S1. Accession numbers for constructed mitogenomes for *Bougainvillea muscus* and *Calvadosia cruxmelitensis* are NCBI Accession ORXXXXX and NCBI Accession ORXXXXX respectively. All code associated with the analyses presented are available in GitHub (https://github.com/ayannadmays/Cnidarian-TE-Analysis). The Medusozoa repeat library generated here will be available in Figshare (forthcoming).

### Competing interests

The authors declare no competing interests.

### Funding

Funding for this work comes from the University of California, Santa Cruz and the Cota-Robles fellowship to ADM.

### Authors’ contributions

AMM and ADM conceived and designed the study. ADM analyzed and interpreted the genomic data regarding repetitive element content and activity. FCC constructed and analyzed mitogenomes and reconstructed the phylogeny presented here. ADM wrote the original manuscript draft. FCC and AMM provided feedback and revisions. All authors read and approved the final manuscript.

## Acknowledgements

We would like to acknowledge Russ Corbett-Detig and Sam Bogan for helpful conversations regarding the approach to the TE analysis. In addition, we thank Rion Parsons for his help with bioinformatics troubleshooting on UCSC’s Hummingbird computing cluster and the members of the Macias-Muñoz lab for their feedback and support.

## Notes

https://github.com/ayannadmays/Cnidarian-TE-Analysis

### Competing Interest Statement

The authors have declared no competing interest.

## References

1. Finnegan DJ. Eukaryotic transposable elements and genome evolution. Trends Genet. 1989 Jan 1;5:103–7. doi:10.1016/0168-9525(89)90039-5

2. Wells JN, Feschotte C. A Field Guide to Eukaryotic Transposable Elements. Annu Rev Genet.2020 Nov 23;54(Volume 54, 2020):539–61. doi:10.1146/annurev-genet-040620-022145

3. Wicker T, Sabot F, Hua-Van A, Bennetzen JL, Capy P, Chalhoub B, et al. A unified classification system for eukaryotic transposable elements. Nat Rev Genet. 2007 Dec;8(12):973–82. doi:10.1038/nrg2165

4. Feiner N. Accumulation of transposable elements in Hox gene clusters during adaptive radiation of Anolis lizards. Proc R Soc B Biol Sci. 2016 Oct 12;283(1840):20161555. doi:10.1098/rspb.2016.1555

5. Ricci M, Peona V, Guichard E, Taccioli C, Boattini A. Transposable Elements Activity is Positively Related to Rate of Speciation in Mammals. J Mol Evol. 2018 Jun 1;86(5):303–10. doi:10.1007/s00239-018-9847-7

6. Kleckner N, Chan RK, Tye BK, Botstein D. Mutagenesis by insertion of a drug-resistance element carrying an inverted repetition. J Mol Biol. 1975 Oct 5;97(4):561–75. doi:10.1016/S0022-2836(75)80059-3

7. Lehrman MA, Schneider WJ, Südhof TC, Brown MS, Goldstein JL, Russell DW. Mutation in LDL Receptor: Alu-Alu Recombination Deletes Exons Encoding Transmembrane and Cytoplasmic Domains. Science. 1985 Jan 11;227(4683):140–6. doi:10.1126/science.3155573

8. Lupski JR. Genomic disorders: structural features of the genome can lead to DNA rearrangements and human disease traits. Trends Genet. 1998 Oct 1;14(10):417–22. doi:10.1016/S0168-9525(98)01555-8 PubMed PMID: 9820031.

9. Robberecht C, Voet T, Esteki MZ, Nowakowska BA, Vermeesch JR. Nonallelic homologous recombination between retrotransposable elements is a driver of de novo unbalanced translocations. Genome Res. 2013 Mar 1;23(3):411–8. doi:10.1101/gr.145631.112 PubMed PMID: 23212949.

10. Assis R, Bachtrog D. Neofunctionalization of young duplicate genes in Drosophila. Proc Natl Acad Sci. 2013 Oct 22;110(43):17409–14. doi:10.1073/pnas.1313759110

11. Ohno S. Evolution by Gene Duplication. Springer Science & Business Media; 2013. 171 p.

12. Teefy BB, Siebert S, Cazet JF, Lin H, Juliano CE. PIWI–piRNA pathway-mediated transposable element repression in Hydra somatic stem cells. RNA. 2020 May 1;26(5):550–63. doi:10.1261/rna.072835.119 PubMed PMID: 32075940.

13. Ying H, Hayward DC, Klimovich A, Bosch TCG, Baldassarre L, Neeman T, et al. The Role of DNA Methylation in Genome Defense in Cnidaria and Other Invertebrates. Mol Biol Evol. 2022 Feb 1;39(2):msac018. doi:10.1093/molbev/msac018

14. Yoder JA, Walsh CP, Bestor TH. Cytosine methylation and the ecology of intragenomic parasites. Trends Genet. 1997 Aug 1;13(8):335–40. doi:10.1016/S0168-9525(97)01181-5 PubMed PMID: 9260521.

15. Cosby RL, Judd J, Zhang R, Zhong A, Garry N, Pritham EJ, et al. Recurrent evolution of vertebrate transcription factors by transposase capture. Science. 2021 Feb 19;371(6531):eabc6405. doi:10.1126/science.abc6405

16. Kazazian HH. Mobile Elements: Drivers of Genome Evolution. Science. 2004 Mar 12;303(5664):1626–32. doi:10.1126/science.1089670

17. Mukherjee K, Moroz LL. Transposon-derived transcription factors across metazoans. Front Cell Dev Biol.2023 Mar 7;11. doi:10.3389/fcell.2023.1113046

18. Park E, Hwang DS, Lee JS, Song JI, Seo TK, Won YJ. Estimation of divergence times in cnidarian evolution based on mitochondrial protein-coding genes and the fossil record. Mol Phylogenet Evol. 2012 Jan 1;62(1):329–45. doi:10.1016/j.ympev.2011.10.008

19. DeBiasse MB, Buckenmeyer A, Macrander J, Babonis LS, Bentlage B, Cartwright P, et al. A Cnidarian Phylogenomic Tree Fitted With Hundreds of 18S Leaves. Bull Soc Syst Biol.2024 Apr 11;3(2). doi:10.18061/bssb.v3i2.9267

20. Kayal E, Roure B, Philippe H, Collins AG, Lavrov DV. Cnidarian phylogenetic relationships as revealed by mitogenomics. BMC Evol Biol. 2013 Jan 9;13(1):5. doi:10.1186/1471-2148-13-5

21. Cartwright P, Nawrocki AM. Character Evolution in Hydrozoa (phylum Cnidaria). Integr Comp Biol. 2010 Sep 1;50(3):456–72. doi:10.1093/icb/icq089

22. Collins AG. Phylogeny of Medusozoa and the evolution of cnidarian life cycles. J Evol Biol. 2002 May 1;15(3):418–32. doi:10.1046/j.1420-9101.2002.00403.x

23. Cunningham CW, Buss LW. Molecular evidence for multiple episodes of paedomorphosis in the family Hydractiniidae. Biochem Syst Ecol. 1993 Jan 1;21(1):57–69. doi:10.1016/0305-1978(93)90009-G

24. Weismann A. Die entstehung der sexualzellen bei den hydromedusen: Zugleich ein beitrag zur kenntniss des baues und der lebenserscheinungen dieser gruppe. G. Fischer; 1883. 426 p.

25. Kon-Nanjo K, Kon T, Horkan HR, Febrimarsa, Steele RE, Cartwright P, et al. Chromosome-level genome assembly of *Hydractinia symbiolongicarpus*. Parker J, editor. G3 Genes Genomes Genet.2023 May 18;jkad107. doi:10.1093/g3journal/jkad107

26. Puzakov MV, Puzakova LV, Shi S, Cheresiz SV. maT and mosquito transposons in cnidarians: evolutionary history and intraspecific differences. Funct Integr Genomics. 2023 Jul 16;23(3):244. doi:10.1007/s10142-023-01175-0

27. Puzakov MV, Puzakova LV. Prevalence, Diversity, and Evolution of L18 (DD37E) Transposons in the Genomes of Cnidarians. Mol Biol. 2022 Jun 1;56(3):424–36. doi:10.1134/S0026893322030104

28. Ulupova YN, Puzakov MV, Puzakova LV. Pogo DNA Transposons in the Genomes of the Aurelia Genus Jellyfish. Mol Genet Microbiol Virol. 2023 Jun 1;38(2):79–85. doi:10.3103/S089141682302009X

29. Wong WY, Simakov O, Bridge DM, Cartwright P, Bellantuono AJ, Kuhn A, et al. Expansion of a single transposable element family is associated with genome-size increase and radiation in the genus Hydra. Proc Natl Acad Sci U S A. 2019 Nov 12;116(46):22915–7. doi:10.1073/pnas.1910106116 PubMed PMID: 31659034; PubMed Central PMCID: PMC6859323.

30. Kon T, Kon-Nanjo K, Simakov O. Subtelomeric repeat expansion in Hydractinia symbiolongicarpus chromosomes. Mob DNA. 2025 Mar 25;16(1):14. doi:10.1186/s13100-025-00355-y

31. Talavera G, Castresana J. Improvement of Phylogenies after Removing Divergent and Ambiguously Aligned Blocks from Protein Sequence Alignments [Internet]. [cited 2026 Apr 16]. Available from: 10.1080/10635150701472164

32. Liu X, Zhao L, Majid M, Huang Y. Orthoptera-TElib: a library of Orthoptera transposable elements for TE annotation. Mob DNA. 2024 Mar 15;15(1):5. doi:10.1186/s13100-024-00316-x

33. Flynn JM, Hubley R, Goubert C, Rosen J, Clark AG, Feschotte C, et al. RepeatModeler2 for automated genomic discovery of transposable element families. Proc Natl Acad Sci. 2020 Apr 28;117(17):9451–7. doi:10.1073/pnas.1921046117

34. Yan H, Bombarely A, Li S. DeepTE: a computational method for de novo classification of transposons with convolutional neural network. Bioinformatics. 2020 Aug 1;36(15):4269–75. doi:10.1093/bioinformatics/btaa519

35. Kimura M. A simple method for estimating evolutionary rates of base substitutions through comparative studies of nucleotide sequences. J Mol Evol. 1980 Jun 1;16(2):111–20. doi:10.1007/BF01731581

36. Cazet JF, Siebert S, Little HM, Bertemes P, Primack AS, Ladurner P, et al. A chromosome-scale epigenetic map of the Hydra genome reveals conserved regulators of cell state. Genome Res. 2023 Feb 1;33(2):283–98. doi:10.1101/gr.277040.122 PubMed PMID: 36639202.

37. Kayal E, Bentlage B, Sabrina Pankey M, Ohdera AH, Medina M, Plachetzki DC, et al. Phylogenomics provides a robust topology of the major cnidarian lineages and insights on the origins of key organismal traits. BMC Evol Biol. 2018 Apr 13;18(1):68. doi:10.1186/s12862-018-1142-0

38. Feng H, Lv S, Li R, Shi J, Wang J, Cao P. Mitochondrial genome comparison reveals the evolution of cnidarians. Ecol Evol. 2023;13(6):e10157. doi:10.1002/ece3.10157

39. Chalopin D, Naville M, Plard F, Galiana D, Volff JN. Comparative Analysis of Transposable Elements Highlights Mobilome Diversity and Evolution in Vertebrates. Genome Biol Evol. 2015 Jan 9;7(2):567–80. doi:10.1093/gbe/evv005 PubMed PMID: 25577199; PubMed Central PMCID: PMC4350176.

40. Chénais B, Caruso A, Hiard S, Casse N. The impact of transposable elements on eukaryotic genomes: From genome size increase to genetic adaptation to stressful environments. Gene. 2012 Nov 1;509(1):7–15. doi:10.1016/j.gene.2012.07.042

41. Kidwell MG. Transposable elements and the evolution of genome size in eukaryotes. Genetica. 2002 May 1;115(1):49–63. doi:10.1023/A:1016072014259

42. Kim HM, Weber JA, Lee N, Park SG, Cho YS, Bhak Y, et al. The genome of the giant Nomura’s jellyfish sheds light on the early evolution of active predation. BMC Biol. 2019 Mar 29;17(1):28. doi:10.1186/s12915-019-0643-7

43. Xia W, Li H, Cheng W, Li H, Mi Y, Gou X, et al. High-Quality Genome Assembly of Chrysaora quinquecirrha Provides Insights Into the Adaptive Evolution of Jellyfish. Front Genet.2020 Jun 4;11. doi:10.3389/fgene.2020.00535

44. Gold DA, Katsuki T, Li Y, Yan X, Regulski M, Ibberson D, et al. The genome of the jellyfish Aurelia and the evolution of animal complexity. Nat Ecol Evol. 2019 Jan;3(1):96–104. doi:10.1038/s41559-018-0719-8

45. Nong W, Cao J, Li Y, Qu Z, Sun J, Swale T, et al. Jellyfish genomes reveal distinct homeobox gene clusters and conservation of small RNA processing. Nat Commun. 2020 Jun 19;11(1):3051. doi:10.1038/s41467-020-16801-9

46. Pascual-Torner M, Carrero D, Pérez-Silva JG, Álvarez-Puente D, Roiz-Valle D, Bretones G, et al. Comparative genomics of mortal and immortal cnidarians unveils novel keys behind rejuvenation. Proc Natl Acad Sci. 2022 Sep 6;119(36):e2118763119. doi:10.1073/pnas.2118763119

47. Santander MD. Evolução genômica dos Medusozoa (Cnidaria), com ênfase em elementos repetitivos nos Scyphozoa.

48. Straehler-Pohl I, Jarms G. Back to the roots, Part 1—early life cycle data of Rhopaliophora (Scyphozoa, Cubozoa and Staurozoa). Plankton Benthos Res. 2022;17(1):1–33. doi:10.3800/pbr.17.1

49. Straehler-Pohl I, Jarms G. Back to the roots, Part 2—Rhopaliophora (Scyphozoa, Cubozoa and Staurozoa) reborn based on early life cycle data. Plankton Benthos Res. 2022;17(2):105–26. doi:10.3800/pbr.17.105

50. Baril T, Pym A, Bass C, Hayward A. Transposon accumulation at xenobiotic gene family loci in aphids. Genome Res. 2023 Oct 1;33(10):1718–33. doi:10.1101/gr.277820.123 PubMed PMID: 37852781.

51. Lyu K, Xiao J, Lyu S, Liu R. Comparative Analysis of Transposable Elements in Strawberry Genomes of Different Ploidy Levels. Int J Mol Sci. 2023 Jan;24(23):23. doi:10.3390/ijms242316935

52. Sotero-Caio CG, Platt RN II, Suh A, Ray DA. Evolution and Diversity of Transposable Elements in Vertebrate Genomes. Genome Biol Evol. 2017 Jan 1;9(1):161–77. doi:10.1093/gbe/evw264

53. Xu Y, Tang Y, Feng W, Yang Y, Cui Z. Comparative Analysis of Transposable Elements Reveals the Diversity of Transposable Elements in Decapoda and Their Effects on Genomic Evolution. Mar Biotechnol. 2023 Dec 1;25(6):1136–46. doi:10.1007/s10126-023-10265-w

54. Durán-Fuentes JA, Maronna MM, Palacios-Gimenez OM, Castillo ER, Ryan JF, Daly M, et al. Repeatome diversity in sea anemone genomics (Cnidaria: Actiniaria) based on the Actiniaria-REPlib library. BMC Genomics. 2025 May 13;26(1):473. doi:10.1186/s12864-025-11591-0

55. Fourreau CJL, Kise H, Santander MD, Pirro S, Maronna MM, Poliseno A, et al. Genome sizes and repeatome evolution in zoantharians (Cnidaria: Hexacorallia: Zoantharia). PeerJ. 2023 Oct 16;11:e16188. doi:10.7717/peerj.16188 PubMed PMID: 37868064; PubMed Central PMCID: PMC10586311.

56. Collins AG. Phylogeny of Medusozoa and the evolution of cnidarian life cycles. J Evol Biol. 2002 May 1;15(3):418–32. doi:10.1046/j.1420-9101.2002.00403.x

57. DeVries DR. The Freshwater Jellyfish Craspedacusta sowerbyi: a Summary of Its Life History, Ecology, and Distribution. J Freshw Ecol. 1992 Mar 1;7(1):7–16. doi:10.1080/02705060.1992.9664665

58. Jankowski T. The freshwater medusae of the world – a taxonomic and systematic literature study with some remarks on other inland water jellyfish. Hydrobiologia. 2001 Oct 1;462(1):91–113. doi:10.1023/A:1013126015171

59. Jankowski T, Collins AG, Campbell R. Global diversity of inland water cnidarians. Hydrobiologia. 2008 Jan 1;595(1):35–40. doi:10.1007/s10750-007-9001-9

60. Khalturin K, Shinzato C, Khalturina M, Hamada M, Fujie M, Koyanagi R, et al. Medusozoan genomes inform the evolution of the jellyfish body plan. Nat Ecol Evol. 2019 May;3(5):811–22. doi:10.1038/s41559-019-0853-y

61. Santander MD, Maronna MM, Ryan JF, Andrade SCS. The state of Medusozoa genomics: current evidence and future challenges. GigaScience. 2022 Jan 1;11:giac036. doi:10.1093/gigascience/giac036

62. Dong Z, Wang F, Liu Y, Li Y, Yu H, Peng S, et al. Genomic and single-cell analyses reveal genetic signatures of swimming pattern and diapause strategy in jellyfish. Nat Commun. 2024 Jul 15;15(1):5936. doi:10.1038/s41467-024-49848-z

63. Matsumoto Y, Piraino S, Miglietta MP. Transcriptome Characterization of Reverse Development in Turritopsis dohrnii (Hydrozoa, Cnidaria). G3 GenesGenomesGenetics. 2019 Dec 1;9(12):4127–38. doi:10.1534/g3.119.400487

64. Miglietta MP. On the perils of working on nonmodel organisms. Proc Natl Acad Sci. 2023 Mar 14;120(11):e2216683120. doi:10.1073/pnas.2216683120

65. Miglietta MP, Piraino S, Kubota S, Schuchert P. Species in the genus Turritopsis (Cnidaria, Hydrozoa): a molecular evaluation. J Zool Syst Evol Res. 2007;45(1):11–9. doi:10.1111/j.1439-0469.2006.00379.x

