## Supplemental figures for "Comparative analysis of transposable elements in jellyfish and hydroid species (Cnidaria: Medusozoa)"

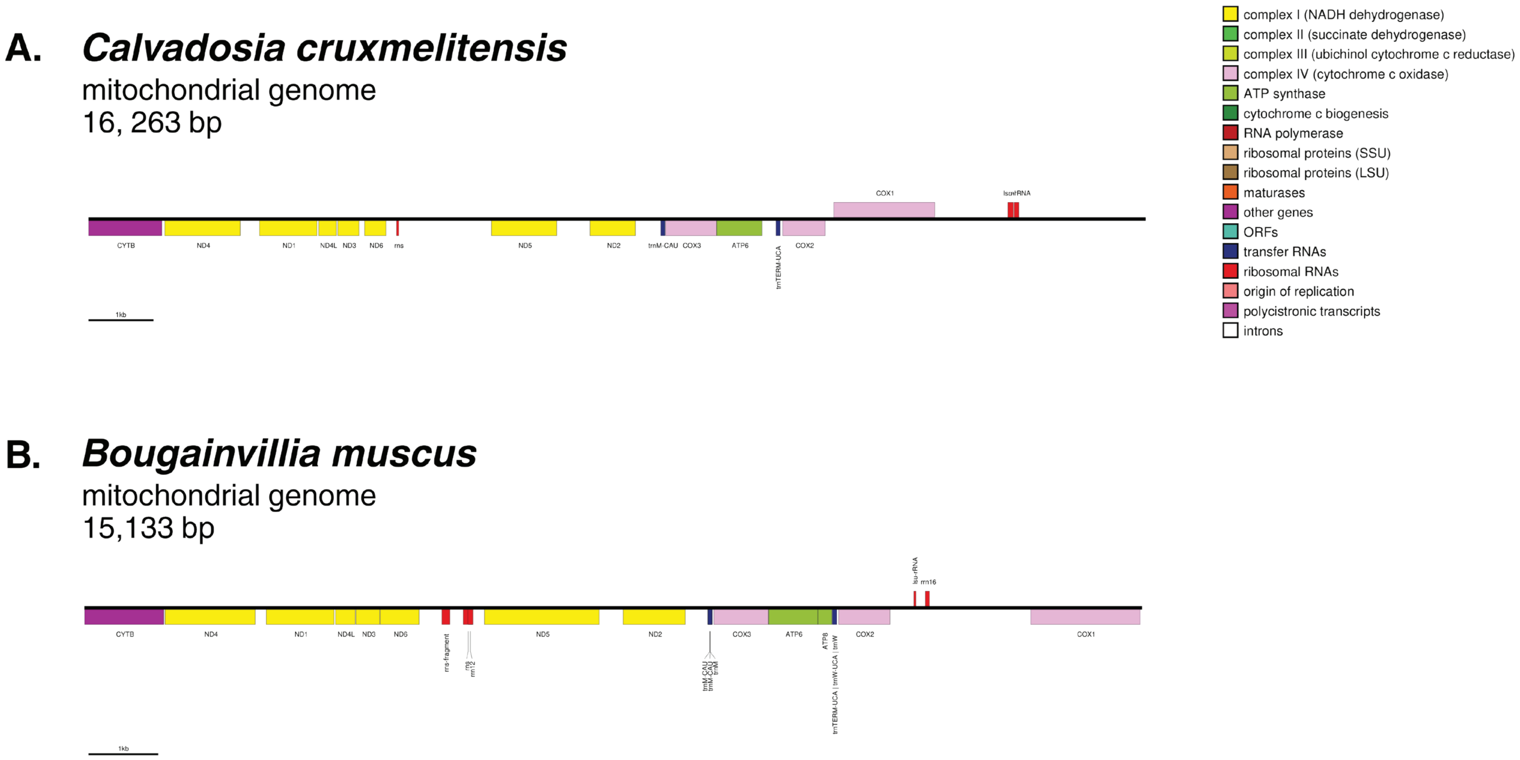


Fig S1. Complete mitochondrial genomes of (A) the staurozoan Calvadosia cruxmelitensis and (B) hyrozoan Bougainvillia muscus. Functional groups are color-coded. Genes transcribed from the heavy and light strands are shown above and below the chain, respectively.

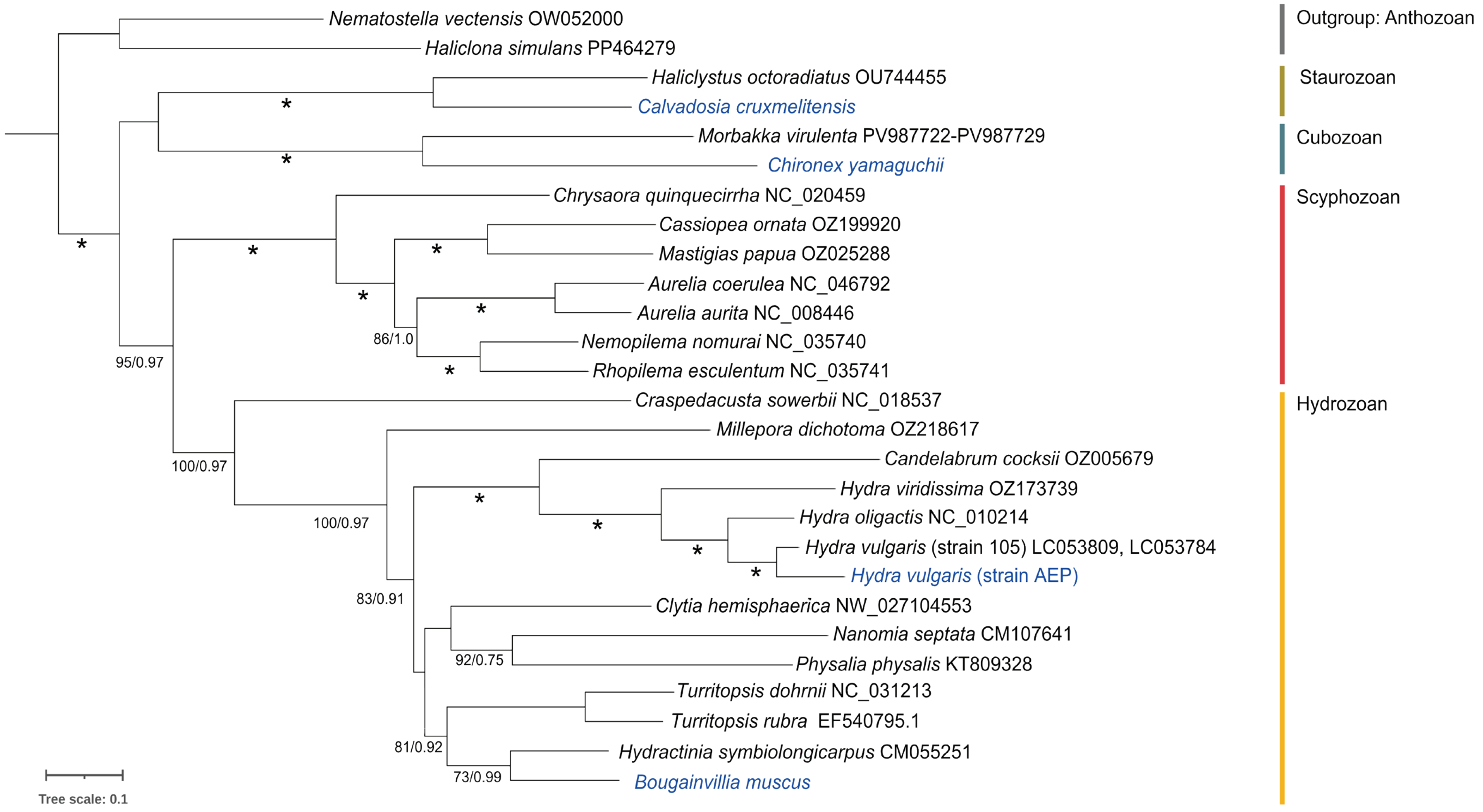


Fig. S2. Maximum likelihood phylogeny inferred in RAxML (GTR + I + G) from 25 medusozoan mitogenomic sequences (12,004 bp; from mitochondrial genes to complete mitogenomes), with two anthozoans as outgroups. Node support values (Maximum Likelihood [ML] / Bayesian Posterior Probabilities [BPP]; ML ≥ 70%, BPP ≥ 0.95) are shown below branches; * indicates full support. Scale bar, substitutions per site. Blue labels denote taxa with partial to near-complete mitochondrial markers newly extracted from published SRA datasets.

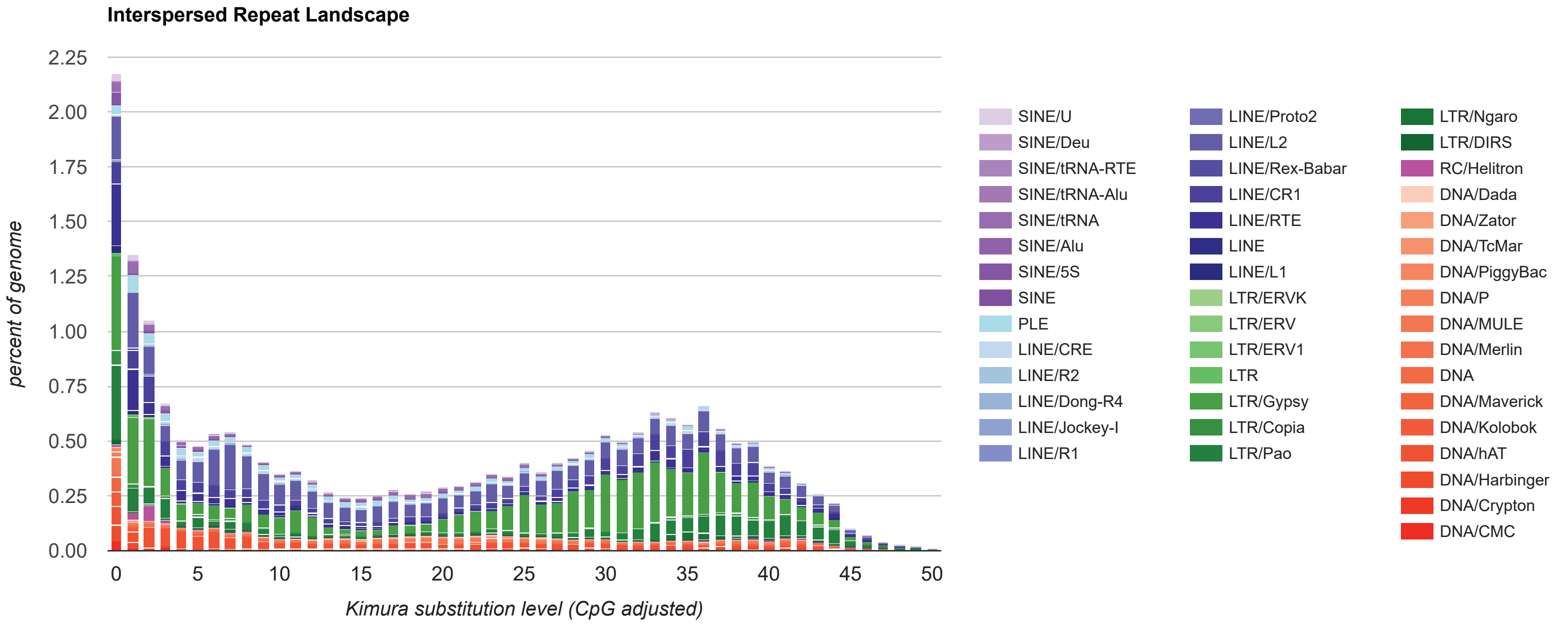


Fig. S3. Repeat landscape of *Haliclystus octoradiatus* generated by using the createLandscape.pl function of RepeatMasker given the .divsum file resulting  rom RepeatMasker with the custom Medusozoa repeat library as input. Inconsistencies in repeat landscapes from both methods precluded its inclusion in analysis.
